# Accurate and Reproducible Functional Maps in 127 Human Cell Types via 2D Genome Segmentation

**DOI:** 10.1101/118752

**Authors:** Yu Zhang, Ross C. Hardison

## Abstract

The Roadmap Epigenomics consortium has published whole-genome functional annotation maps in 127 human cell types and cancer cell lines by integrating data from multiple epigenetic marks. These maps have thereby been widely used by the community for studying gene regulation in cell type specific contexts and predicting functional impacts of DNA mutations on disease. Here, we present a new map of functional elements produced by a recently published method called IDEAS on the same data set. The IDEAS method has several unique advantages and was shown to outperform existing methods, including the one used by the Roadmap Epigenomics consortium. We further introduce a simple but highly effective pipeline to greatly improve the reproducibility of functional annotation. Using five categories of independent experimental results, we extensively compared the annotation produced by IDEAS and the Roadmap Epigenomics consortium. While the overall concordance between the two maps was high, we observed many differences in the details and in the position-wise consistency of annotation across cell types. We show that the IDEAS annotation was uniformly and often substantially more accurate than the Roadmap Epigenomics result. This study therefore reports on the quality of an existing functional map in 127 human genomes and provides an alternative and better map to be used by the community. The annotation result can be visualized in the UCSC genome browser via the hub at http://bx.psu.edu/~yuzhang/Roadmap_ideas/ideas_hub.txt

## INTRODUCTION

Thousands of epigenetics data sets have been released in hundreds of human cell types (ENCODE Project Consortium, 2012; Roadmap Epigenomics Consortium, 2015; Stunnenberg et al. 2016), which is an incredibly rich resource of information for studying epigenetic events towards understanding gene regulation in the human genome. The raw data generated by high-throughput sequencing technologies are however difficult to interpret, as not only the signals are noisy, but also different epigenetic marks may represent distinct regulatory functions in a combinatorial fashion. To facilitate the discovery and interpretation of functional elements in human genomes, computational algorithms such as genome segmentation (Ernst and Kellis, 2012; Hoffman et al., 2012) have been developed to annotate the human genome based on multiple epigenetic data sets. The principle is to identify *de novo* combinatorial patterns of multiple epigenetic marks, which are called epigenetic states, within intervals across the genome. The epigenetic states inferred by genome segmentation methods have been experimentally shown to correspond to unique functional elements and have impacts on phenotypes (Hardison, 2012). Epigenetic states are a low-dimensional de-noised representation of the high-dimensional raw data, and thus are more convenient for visualization, interpretation and testing in downstream analyses.

Currently the Roadmap Epigenomics project has published its own genome segmentation results in 111 human cell types and 16 cell types from the ENCODE project including a few cancer cell lines (Roadmap Epigenomics Consortium, 2015). These results have thereby been widely used by recent studies to facilitate the discovery and interpretation of new biological insights, especially for interpreting and prioritizing non-coding variants in human complex diseases (Pickrell, 2014; Chung et al., 2014; Kichaev et al., 2014; Kichaev and Pasaniuc, 2015; Farh et al., 2015; Li and Kellis, 2016; Lu et al., 2016). The Roadmap segmentations were produced from a popular algorithm called ChromHMM (Ernst and Kellis, 2012), which employs a Hidden Markov model with binary emission probability to identify epigenetic states. The algorithm works by first converting the raw signals in 200bp windows to binary values based on a significance cutoff in each data set, and then concatenating the epigenomes of all cell types together in a linear fashion for joint segmentation. An advantage of their approach is the computational speed and simple interpretation of the result, as the method deals with binary outcomes and only models 1-dim data dependence across the genome. The disadvantage of ChromHMM is however significant. First, by converting quantitative data to binary values, magnitude of the signals will be lost and the results will be subject to threshold choices. Secondly, the number of epigenetic states must be predetermined, which is error prone and may miss important epigenetic states. Thirdly and more importantly, ChromHMM does not account for the fact that all cell types share the same underlying DNA sequences and hence the regulatory events across cell types are not independent. ChromHMM is thus a “1D” segmentation method not optimally designed for joint segmentation of multiple epigenomes.

We recently introduced a new genome segmentation algorithm called IDEAS (Zhang et al., 2016) that tackled the above issues. IDEAS works on quantitative data without binarization, although binary data can always be used as a special case. IDEAS employs Bayesian non-parametric techniques to automatically choose the number of states from the data instead of requiring user input. However the user can still fix the number of states whenever desired. Importantly, IDEAS is a “2D” segmentation method that, in addition to modeling data dependence along the genome, further accounts for the position-wise correlation between regulatory events across different cell types. Inference of the IDEAS model has a linear time complexity with respect to the genome size and the number of cell types involved. The method is thus computationally efficient even for analyzing hundreds of cell types simultaneously. Our previous studies have shown that IDEAS could run as fast as ChromHMM.

In light of the advantages of the IDEAS method over existing genome segmentation tools, we introduce a new map of regulatory elements in the 127 Roadmap epigenomes generated by the IDEAS method. We used the same five histone marks as used by ChromHMM to produce the map, because they are available in all of the 127 cell types. Here, we present a comprehensive evaluation of the results produced by IDEAS and ChromHMM by using fived independent sources of experimental data sets, including the RNA-seq data in 56 cell types from the Roadmap Epigenomics project, the expression quantitative trait loci (eQTL) in 44 tissues from the GTEx project (GTEx consortium, 2015), the enhancer usage data in 808 human CAGE libraries from the FANTOM5 project (Andersson et al. 2014), four sequence-based scores for functional impact of DNA mutations (Davydov et al., 2010; Kircher et al., 2014; Gulko et al., 2015; Maurato et al., 2015), and the promoter capture Hi-C data in 17 blood cell types from the IHEC project (Javierre et al., 2016). These data sets are independent of the data sets used for generating the functional maps. Correlation with these experimental results could therefore reflect the accuracy of the two methods with respect to predicting both functional potentials and chromatin structural information using epigenetic states.

## RESULTS

### Joint segmentation in 127 epigenomes

We ran IDEAS on the uniformly processed p-value tracks of five histone marks (H3K4me3, H3K4me1, H3K36me3, H3K27me3, H3K9me3) commonly available in the 127 epigenomes. Our program automatically identified 20 epigenetic states from the quantitative signals. Comparing the mean signals of our epigenetic states with those of the 15 states by ChromHMM on the same data, we found most of the fundamental states were commonly identified with similar proportions between the two results, such as active transcription start sites (TssA), enhancers (Enh), bivalent TSS (TssBiv) and bivalent enhancers (Enh Biv), heterochromatin (Het), repressed polycomb (ReprPC) and quiescent regions (Quies) (Figure 1a). For consistency, we adopted the mnemonics used by Roadmap Epigenomics consortium on the 15-state model to assign labels to our states. A brief interpretation of our mnemonics assignment is given in Table S1. In addition, our states captured some novel patterns in the quantitative signals of epigenetic marks and their combinations, as shown by several novel states carrying combinatorial patterns of moderate enhancers, heterochromatin and repressive marks.

**Figure 1.**
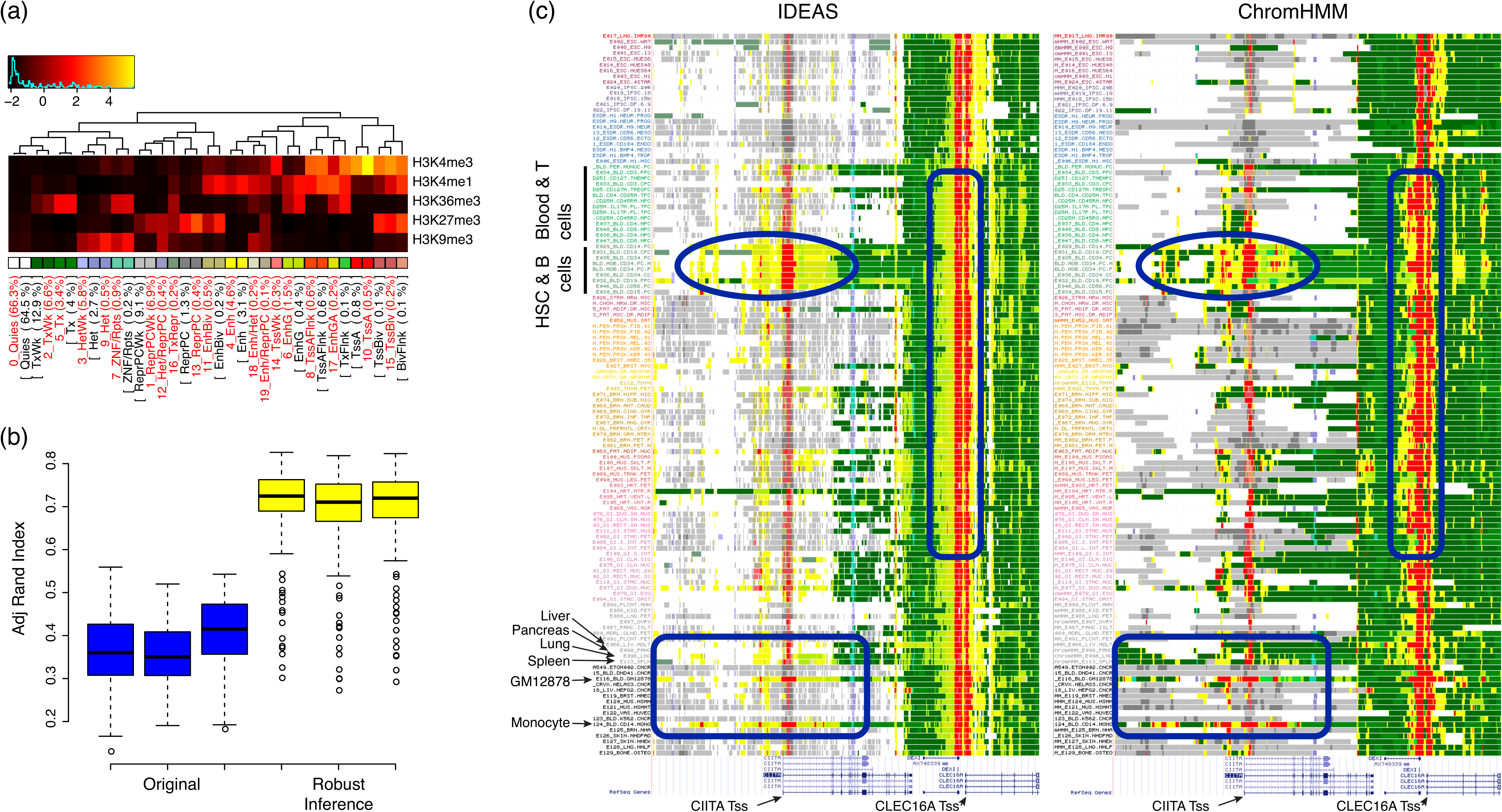
Annotation of functional elements in the 127 Roadmap cell types. (a) Mean epigenetic signal in the IDEAS inferred states (red labeled) and the ChromHMM inferred states (black labeled in brackets). Color key for each epigenetic state is shown below the heatmap. (b) Reproducibility of segmentation by IDEAS between three independent runs using the original program (blue) and the proposed training pipeline (yellow). Each box shows the agreement measured by adjusted rand index between all pairs of cell types between runs. (c) Example of annotation results by IDEAS and ChromHMM in the 127 cell types at genes *CIITA* and *CLEC16A.* Blue boxes highlight some differences between the two maps. Color keys of epigenetic states are given in (a).

While we do not know if these computationally predicted *de novo* states denote unique biological functions, they were consistently recaptured in different runs of IDEAS. Here, we developed a novel training pipeline (see Methods) of IDEAS that guaranteed to generate reproducible epigenetic states. Briefly, we first performed minibatch training of the IDEAS model to generate a collection of epigenetic states. We then used these states to evaluate reproducibility and consolidate the states into a set of reproducible states. This simple pipeline empirically generated highly reproducible results between independent runs (Figure 1b), and hence the novel states we identified in this study were largely robust.

By visualizing our 20-state model in the UCSC genome browser and comparing with the Roadmap Epigenomics 15-state model, we observed, as expected, substantial agreement between the two maps at a large scale but with some substantial differences in the details. For example, at the *CIITA* and *CLEC16A* genes (Figure 1c), our map was more fine-grained, and at most positions, the state assignments matched across cell types more than in the ChromHMM map. On the other hand, notably different state assignments were observed between the two methods for enhancer, TSS and transcription states. For instance, ChromHMM had many more TSS states (red color) assigned to positions away from known TSS than IDEAS did. ChromHMM annotated transcription states (green color) in liver, pancreas, lung and spleen both within the *CIITA* gene and its upstream non-coding regions. In contrast, IDEAS only annotated transcription states and enhancer states with transcription marks in lung and spleen within the *CIITA* gene, whereas liver and pancreas only had transcription states assigned towards the end. This greater cell type-specificity in expression inferred from the IDEAS segmentation was confirmed by examining independent gene expression data from both Roadmap Epigenomics and GTEx projects, as both showed that *CIITA* was expressed in lung and spleen, but little in liver and pancreas.

### Correlation with RNA-seq

We used RNA-seq data from Roadmap Epigenomics in 56 cell types to evaluate the accuracy of the annotations by IDEAS to predict RNA levels across cell types. We used a functional regression model (Ramsay and Silverman, 1997) to include all epigenetics states within ±110kb of each gene as predictors, where we assumed that the effects of epigenetic states on expression is a smooth curve with respect to their distances to genes. As shown in Figure 2a, while the epigenetic states by both methods were highly predictive of gene expression, the states by IDEAS had a consistently and significantly better power in all cell types (p-value 7.6e-8).

**Figure 2.**
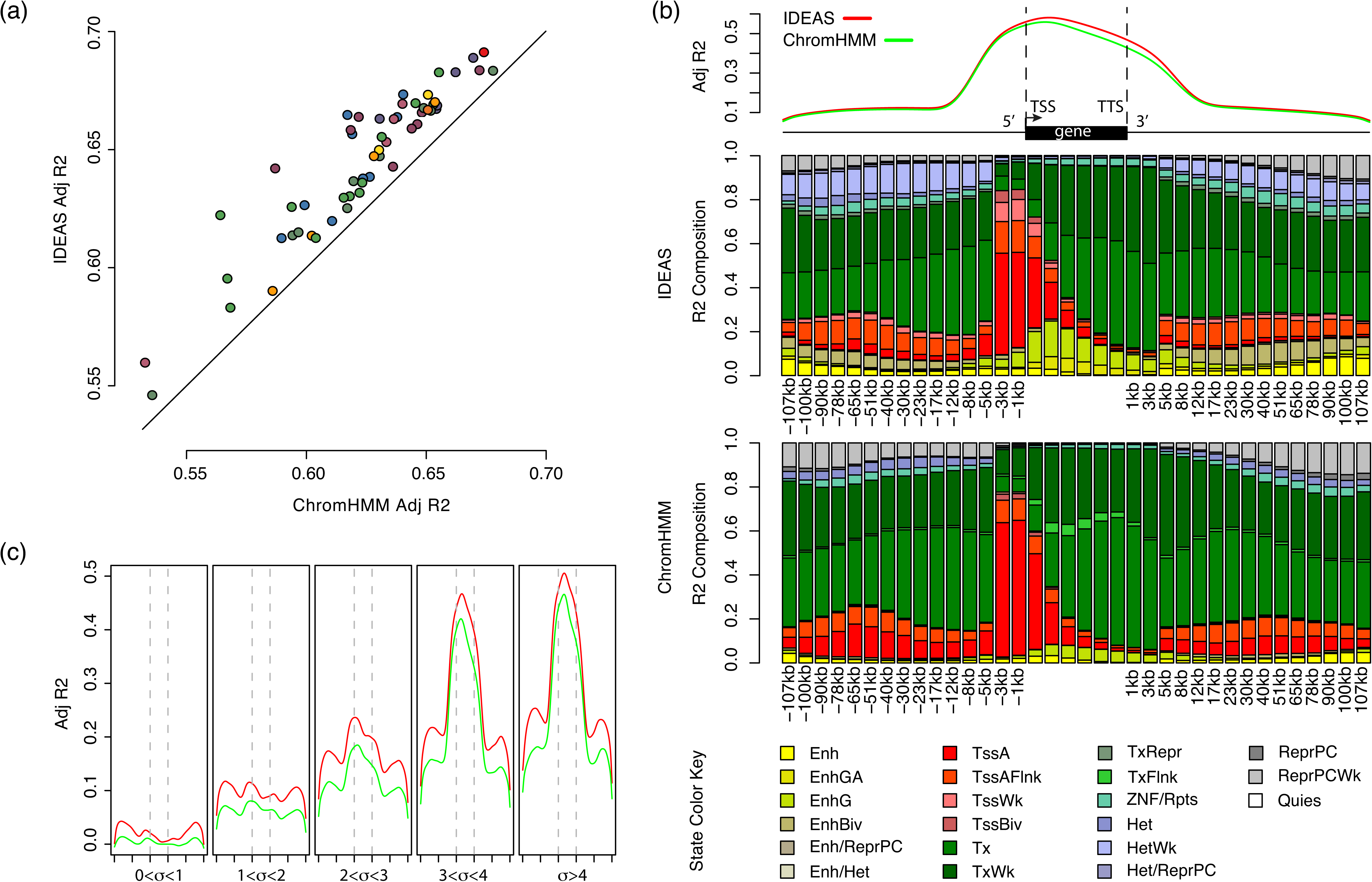
Gene expression explained by epigenetic states. (a) Within-cell type prediction of gene expression in 56 Roadmap cell types. Each point shows the result of one cell type, color-coded by those defined by the Roadmap Epigenomics consortium. (b) Epigenetic state contribution to gene expression as a function of distance to genes. The barplots show the individual state contribution to expression. Color keys for states are shown at the bottom. (c) Prediction of differential gene expression across the 56 cell types. The genes are stratified into five groups by their expression standard deviation. Each panel shows the expression prediction by epigenetic states as a function of distance to gene (x-axis, in the same scale as in (b), and the two vertical dashed lines in each panel show the TSS and TTS locations, respectively). Red: IDEAS; green: ChromHMM.

Investigating the contribution of individual states to expression, as a function of distance to genes, showed that the main difference in prediction power between the two methods occurred towards and beyond transcription termination sites (TTS) of genes (Figure 2b). As expected, we observed predominant contributions of promoter-like states (TssA, TssAFlnk) near the TSS of genes, and transcription states (Tx, TxWk) throughout (Figure 2b). We observed stronger contributions of the genic enhancers (EnhG) within genes and other enhancer states (Enh, EnhBiv) before TSS and after TTS by the IDEAS annotation. The effects of epigenetic states on expression were consistently positive or negative at all distances to genes for both methods, but the effect sizes varied (Figure S1).

Orthogonal to the within-cell type prediction, we next compared the power for predicting differential expression across cell types between the two methods. Within all groups of genes stratified by the levels of differential expression, IDEAS consistently outperformed ChromHMM (Figure 2c). We observed three peaks of adjusted *r^2^* values, within genes near TSS, 50kb upstream of TSS and 50kb downstream of TTS. These three peaks were produced by both methods. The peaks could be attributed to the annotated promoter and enhancer states near the genes, and they also could reflect, in part, the activities of regulatory elements in neighboring genes.

### Prediction of FANTOM5 enhancers

We evaluated the power of epigenetic states for predicting enhancers. We calculated the correlation between the epigenetic states and the FANTOM5 enhancer usage data by a linear regression model. In the FANTOM5 project, enhancer RNA data derived from CAGE libraries from a series of 808 normal tissues, cell types and cancer cell lines was used to estimate enhancer usage in each cell type. We calculated the correlation between each pair of CAGE library and Roadmap cell type separately. As shown in Figure 3a, in comparisons with the ChromHMM annotations, the annotations from IDEAS were significantly better correlated with the FANTOM5 enhancer usage data in almost all CAGE libraries. On average, our map had 25% more power (in terms of adjusted *r^2^*) than the ChromHMM map for predicting enhancer usage. Further investigation of the relative patterns of prediction in all pairs of CAGE library and Roadmap cell type revealed cell type-specific prediction (Figure 3b), such as enhancers in blood, brain, and epithelial cell types. These results confirmed the power of epigenetic states for predicting cell type-specific enhancers.

**Figure 3.**
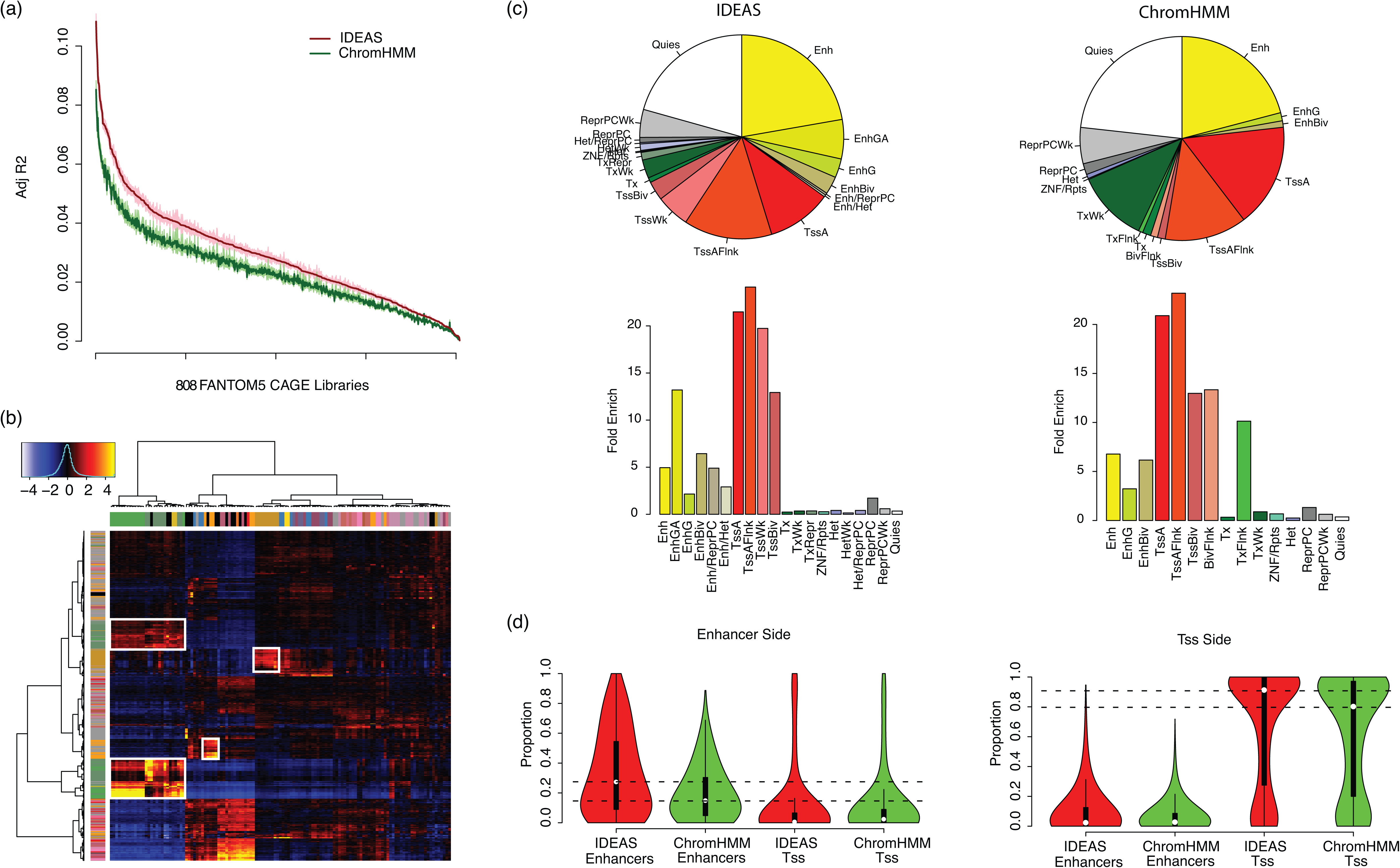
Accuracy for predicting FANTOM5 enhancers. (a) Prediction of tissue specific enhancers in 808 FANTOM5 CAGE libraries. The dark lines show the mean adjusted *r^2^* of predictions by the 127 Roadmap cell types, and the shaded area shows the 95% confidence intervals of means. (b) Z-scores of the adjusted *r^2^* values of tissue-specific enhancer predictions, calculated by removing row and column means and dividing by the overall standard deviation. Cell type specific predictions (similar cell types between Roadmap and FANTOM5) are highlighted in boxes, such as blood cell types (the two boxes on the left), brain tissues (the box in the middle) and epithelial cells (the box on the right). (c) State composition and enrichment within the significant FANTOM5 enhancers, averaged over all Roadmap cell types and CAGE libraries. The enrichment was calculated against genome average. (d) Composition of enhancer and TSS related states in the FANTOM5 significant enhancer-TSS interacting regions.

The state compositions within the significant FANTOM5 enhancer peaks (averaged over 808 CAGE libraries and 127 Roadmap Epigenomes) were notably different between the two methods (Figure 3c). About 68% of the enhancer peak regions were annotated as either enhancer or promoter related states by IDEAS, whereas ChromHMM assigned only 54% of the regions to similar states, with an additional proportion of the regions annotated as weak transcription (TxWk). Notably, the enhancer states with transcription marks (EnhG, EnhGA) by IDEAS had a substantially larger proportion than ChromHMM. Further calculation of fold enrichments showed that the epigenetic states were similarly enriched in the enhancer peak regions by both methods. Taken together, the results of these comparisons showed that annotations generated by the IDEAS segmentations were better predictors of FANTOM5 enhancers.

The FANTOM5 Consortium has identified ~56,000 significant enhancer-TSS pairs showing correlated regulatory activities across the CAGE libraries. We investigated whether the epigenetic states and their pairing were enriched within and between the enhancer-TSS regions. The states generated by both IDEAS and ChromHMM showed a substantial enrichment of enhancer and TSS states assigned in the enhancer-TSS regions (Figure 3d). However, IDEAS annotated more enhancer states in the enhancer side of the paired regions, and more TSS states in the TSS side of the paired regions. By accounting for the marginal enrichment of states in these regions, we further identified some pairwise combinations of states that were enriched or depleted between the enhancer-TSS regions (Figure S2). Consistent with expectations, enhancer states were frequently paired with TSS states, repressed states tended to pair with low or repressed states, and the enrichment pattern of state pairs depended on gene expression (Figure S3). These results demonstrated that the pairing of the epigenetic states between functionally correlated remote regions were predictive of potential trans-regulations.

### Prediction of eQTLs

Several studies have shown that eQTLs are significantly enriched in regulatory elements (Nicolae et al., 2010; Zhong et al., 2010; GTEx Consortium, 2015). Correlation between epigenetic states and eQTLs therefore could be used to assess annotation accuracy. We analyzed the significant eQTLs (nominal p-val <1e-5) in the 44 tissues from the GTEx project for their correlations with epigenetic states. Due to linkage disequilibrium, most of the eQTLs are likely non-causal, but rather linked to causal single nucleotide polymorphisms (SNP). We therefore clustered nearby eQTLs together as an eQTL interval, and calculated a weighted composition of epigenetic states within each eQTL interval, where the weights are positively correlated with the significance of eQTLs and negatively correlated with the distance to eQTLs within the interval. We further calculated an inversely weighted composition of epigenetic states within the same eQTL intervals as controls. In this way, the genomic background was the same in both cases and controls. We then used logistic regression to predict eQTLs intervals against controls in each GTEx tissue by each Roadmap cell type. As shown in Figure 4a, the epigenetic states by both IDEAS and ChromHMM are predictive of eQTLs, but IDEAS significantly outperformed ChromHMM in all tissues. Further analysis of state enrichment in the eQTL intervals relative to the controls showed that the Tss-related states were most strongly enriched in eQTLs, followed by transcription and enhancer related states, and the heterochromatin states and repressed polycomb states were depleted in eQTLs (Figure 4b, Figure S4).

**Figure 4.**
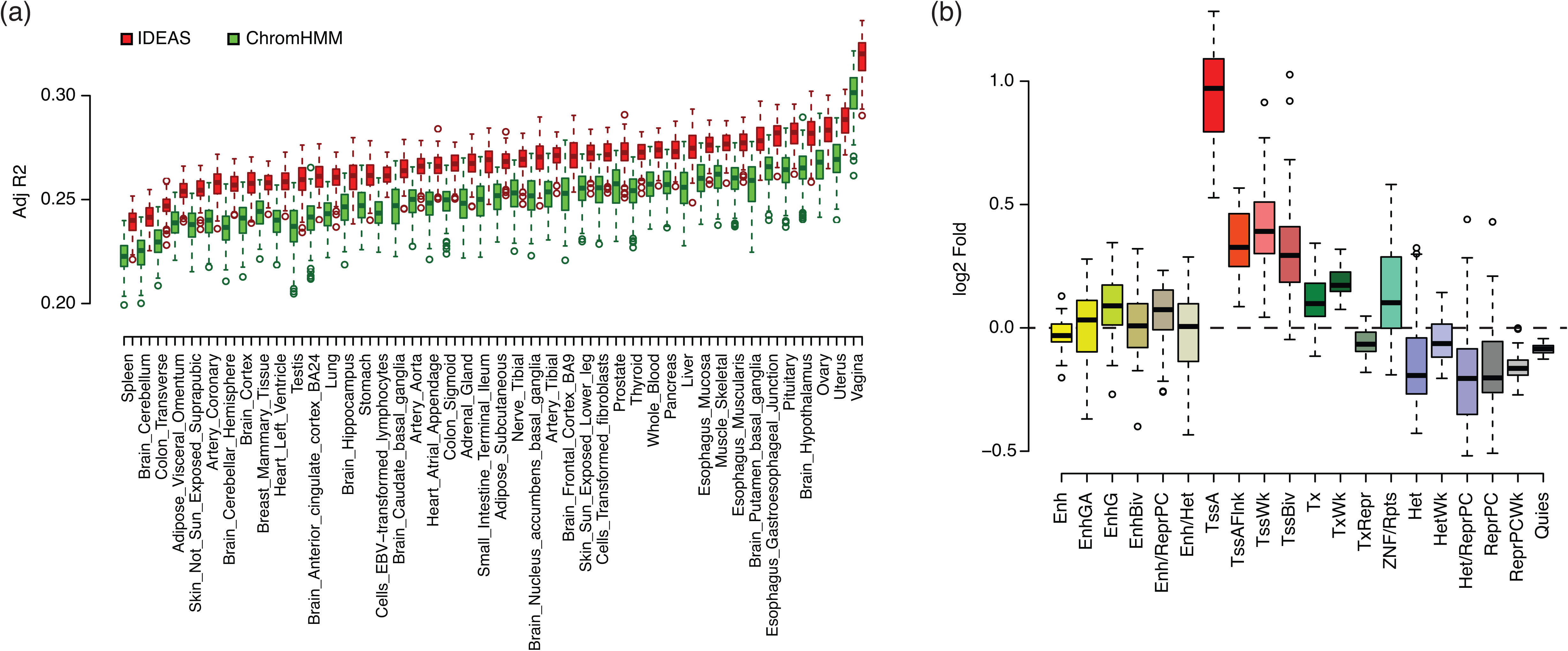
Prediction of eQTLs and state enrichment. (a) Adjusted *r^2^* for predicting eQTLs in 44 GTEx tissues. Each box contains the adjusted *r^2^* values fitted by the epigenetic states in each of the 127 cell types. (b) Enrichment of epigenetic states in eQTLs relative to controls by the IDEAS method. Each box shows the enrichments of the state in the 127 cell types. The color key of epigenetic states is the same as those used in previous figures.

### Correlation with sequence-based functional scores

Numerous sequence-based scores for predicting function of nucleotides or polymorphisms have been computed in the human genome. We included four scores in this study: 1) the Genomic Evolutionary Rate Profiling (GERP) score (Davydov et al., 2010), which identifies functionally constrained elements in multiple alignments; 2) the Combined Annotation Dependent Depletion (CADD) score (Kircher et al., 2014), which predicts deleterious effects of DNA mutations; 3) the fitness consequence of functional annotation (FitCons) score (Gulko et al., 2015), which integrates functional assays with selective pressure to score fraction of genomic positions evincing a pattern of functional assays that are under selection; and 4) the Contextual Analysis of TF Occupancy (CATO) score (Maurano et al., 2015), which quantifies effects of point mutations on transcription factor binding *in vivo.* Using these pre-computed scores for genome-wide mutations, we could assess how useful epigenetic state annotations will be for predicting and interpreting functional impacts of noncoding variants. These scores could also be used to evaluate the positional precision of our annotations, as the scores were calculated at higher resolutions than our annotation maps. Using linear regression on the log-transformed scores, we observed that the states generated by IDEAS significantly and substantially better predicted all scores than did the states generated by ChromHMM (Figure 5a). As we shifted the scores away from their original positions, the prediction by both methods dropped quickly. These results thereby demonstrated that the IDEAS annotation has a better positional precision for predicting functional elements than ChromHMM.

**Figure 5.**
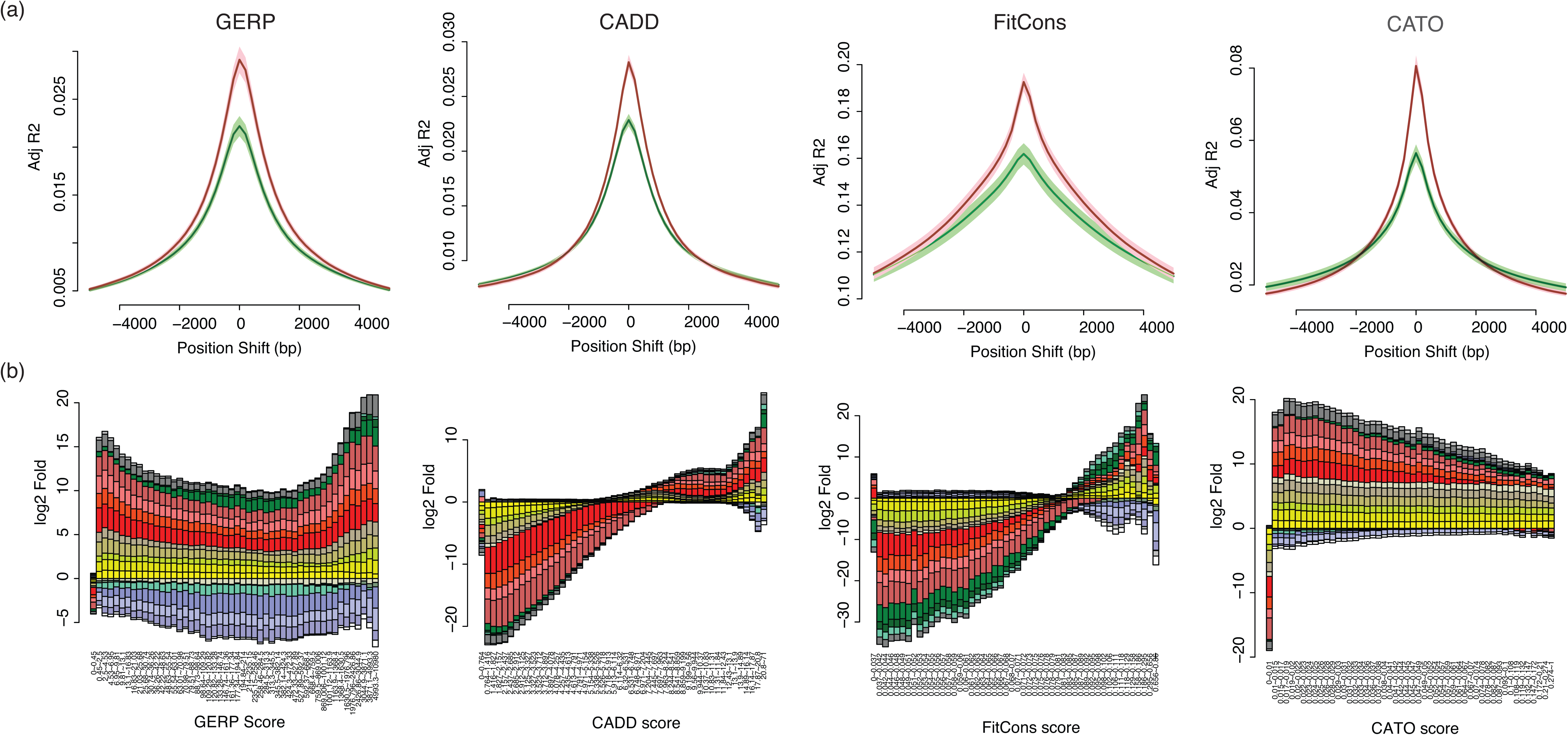
Correlation with sequence-based functional scores and positional precision. (a) Score prediction comparison by the epigenetic states of IDEAS (red) and ChromHMM (green). The positions of scores are shifted to evaluate positional precision of annotations. (b) Cumulative enrichment of epigenetic states by IDEAS as a function of scores. The enrichment was calculated against the genome-wide state distribution. State depletions (negative values) and state enrichments (positive values) are shown in log2 scale. The color key of epigenetic states is the same as those used in previous figures.

The enrichment patterns of epigenetic states with respect to the scores were different for the four scores (Figure 5b), but were relatively consistent between the two methods (Figure S5). In general, the active epigenetic states such as enhancer-, Tss-and transcription-related states were enriched in higher scores, and the inactive epigenetic states such as heterochromatin and quiescent states were enriched in lower scores. Interestingly, the repressed polycomb states (ReprPC, ReprPCWk) were also slightly but consistently enriched in higher scores. This likely resulted from the fact that we calculated the state enrichment using all cell types combined, and the repressed polycomb states often co-occurred with the bivalent TSS and enhancer states (TssBiv, EnhBiv) at the same loci in different cell types.

### Evaluation by promoter-capture HiC

We finally used the promoter-capture HiC data in 17 blood cell types from the IHEC project (Javierre, et al., 2016) to evaluate the ability of our annotations for predicting chromatin structures. We used the states within both the bait and the captured regions to predict the CHiCAGO (Capture HiC Analysis of Genomic Organisation) interaction scores (Cairns et al., 2016). As we have shown in the FANTOM5 data, the state cooccurrence between the two interacting regions were nonrandom. This was also true in the promoter-capture HiC data. We found that including pairwise interaction terms between the states in the bait and the target regions in regression consistently outperformed the corresponding additive models (Figure S6). As a result, we used the interaction model to predict the CHiCAGO interaction scores in each of the 17 IHEC blood cell types using epigenetic states in each of the 127 Roadmap cell types. As shown in Figure 6a, IDEAS uniformly better predicted the CHiCAGO scores in all of the 127 Roadmap cell types, and in general, the Roadmap blood cell types (Blood & T cells and HSC & B cells) had the best prediction power. Also, within each of the 17 IHEC blood cell types, the epigenetic states by IDEAS in the Blood & T and HSC & B cells consistently outperformed ChromHMM (Figure 6b).

**Figure 6.**
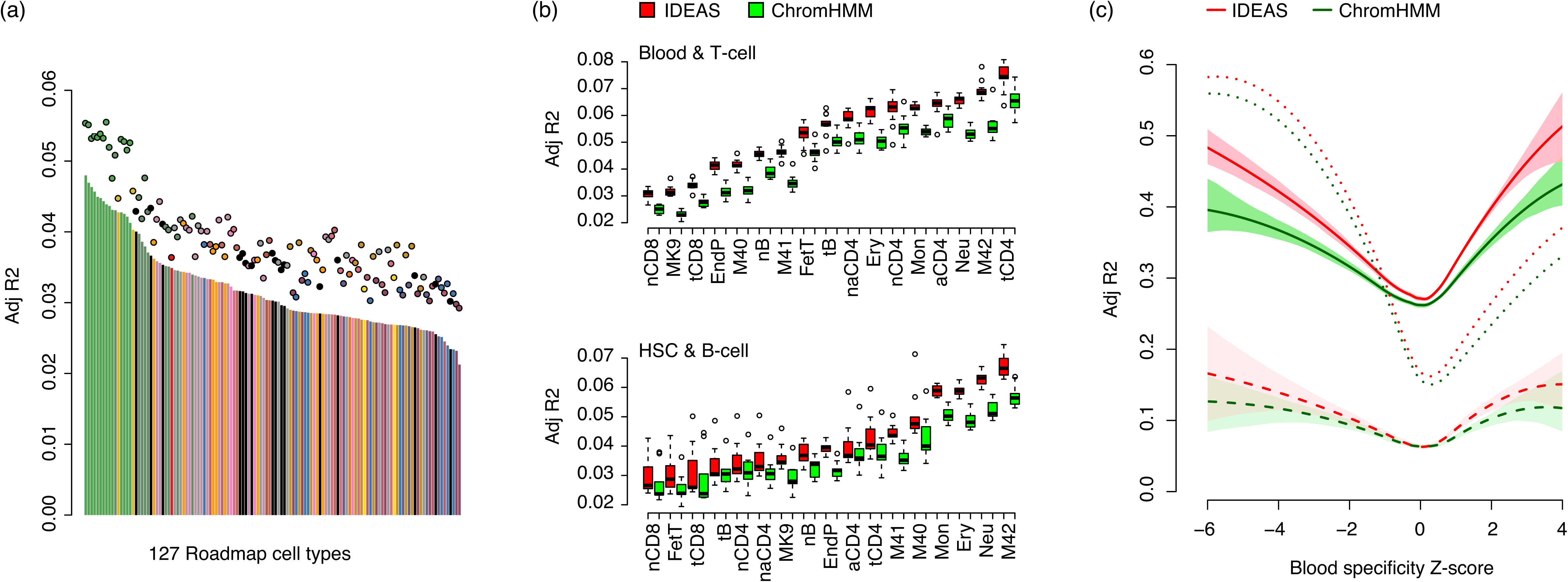
Prediction of chromatin contact in blood cell types. (a) State correlation with CHiCAGO interaction scores in the 127 cell types. Bars show the result by ChromHMM states, and dots show the results by IDEAS states. Cell type colors are assigned by the Roadmap Epigenomics Consortium, where green and dark green indicates Blood & T cells and HSC & B cells, respectively. (b) Comparison of adjusted *r^2^* values in each of the 17 IHEC blood cell types. Red: IDEAS; green: ChromHMM. (c) Prediction of bait gene expression by the epigenetic states in the bait and the captured regions as a function of blood cell type specificity (z-scores, x-axis). Dashed lines: mean adjusted *r^2^* of bait gene expression explained by the states within individual captured regions. Solid lines: mean adjusted *r^2^* of bait expression explained by the sum of states in all captured regions for the same bait. Dotted lines: mean adjusted *r^2^* of bait expression explained by the states within bait regions. Shaded areas show the 95% confidence interval of means.

Finally, we used RNA-seq data to evaluate if the promoter-captured regions carry functional elements that may impact gene expression. As shown in Figure 6c, the epigenetic states by both methods within individual promoter-captured regions were in general correlated with the expression of the bait genes. The correlation however was not strong. After we added the states in all promoter-captured regions together for the same bait, we observed a much stronger correlation with expression. In addition, the epigenetic states of both methods at the bait regions were also strongly correlated with the expression of the bait gene. In all cases, the correlation was stronger for genes up or down regulated specifically in blood cell types. Comparing between the two methods, the IDEAS states showed an overall greater correlation with expression than the ChromHMM states.

## DISCUSSION

We have presented a new functional annotation map jointly produced in the 127 human cell types. Using various independent experimental results, we observed that the epigenetic states of both IDEAS and ChromHMM were useful for predicting functional and structural information of the genome. Our comparative results further showed that the IDEAS map had notable and significant improvement over the ChromHMM map for predicting regulatory events both within and across cell types. At each genomic position, the IDEAS map shows more consistency in epigenetic state annotation across cell types. Simultaneously, it better captured epigenetic variation across cell types as reflected by correlation with differential gene expression.

An important issue in genome segmentation that we attempted to tackle in this study is reproducibility. Annotations produced by the same method under the same parameter settings must be concordant between independent runs in order to be useful. This is a notoriously challenging problem, as no global optimum is guaranteed. Our experience with running existing genome segmentation tools showed that the maps could vary substantially between runs simply by chance. We therefore developed an intuitive, simple and effective approach that substantially improved the reproducibility of our maps. The IDEAS map presented here thus offers an alternative, reliable and more accurate annotation of functional elements in a wealth of human cell types for the community.

The functional map presented in this study has some limitations. First, we only used five core histone marks available in all Roadmap cell types to produce the map, which are ideal for detecting basic functional elements such as enhancers, promoters and repressive states. They do not however have sufficient power to capture more specific regulatory elements such as insulators or occupancy by transcription factors. The Roadmap Epigenomics project has released additional maps using more epigenetic marks either in subsets of cell types or in all cell types via data imputation (Roadmap Epigenomics Consortium, 2015). We have yet to include those additional marks in this study. Secondly, the models used in this study for correlating epigenetic states with the independent validation data were mostly linear. While the maps could possibly be related with other data in non-linear ways, linear models offer simple interpretation of the results. Thirdly, interpreting the inferred epigenetic states, assignment of mnemonics, and state visualization remain challenging problems, particularly when the number of states grows large. In this study we adopted the mnemonics used in Roadmap Epigenomics, which could introduce errors and bias. It would be more desirable to develop learning algorithms for *de novo* interpretation and visualization of the epigenetic states automatically.

Beyond generating functional maps, there are several potential applications uniquely enabled by our 2D segmentation approach. First, the method can naturally leverage information from existing annotations in published cell types to detect functional elements in new cell types and experimental conditions. Our modeling of data dependence across cell types is unsupervised and local in the genome, such that even heterogeneous cell types may be annotated using existing results without cell type matching. Secondly, our joint modeling approach can easily accommodate missing epigenetic marks. New cell types with just one or two epigenetic mark can still be annotated and benefit from the full spectrum of information provided by all epigenetic marks in the published results. This strategy does not require data imputation, and hence is devoid of imputation bias and can save substantially on computing time and disk storage. Thirdly, functional maps produced in the genome of one species may be lifted over to other species in the conserved DNA sequences. Data sets generated in different species may also be integrated and compared via 2D modeling. Towards this end, we have lifted over the map in this study to the mouse genome in mm10 (http://bx.psu.edu/~yuzhang/Roadmap_ideas/mm10_hub.txt), which will be useful as functional elements are largely conserved between human and mouse at the conserved DNA sequences (Xiao et al., 2012). Finally, we note that our method can in general be used to annotate any entities of subjects in a broader scope of gene regulation studies, such as different cell types, experimental conditions, individuals, species and time points.

## MATERIALS AND METHODS

### Roadmap epigenetic data sets

We downloaded the log p-value tracks of a core set of 5 chromatin marks (H3K4me3, H3K4me1, H3K36me3, H3K27me3, H3K9me3) assayed in all of the 127 epigenomes from http://egg2.wustl.edu/roadmap/data/byFileType/signal/consolidated/macs2signal/pval/. We processed the signal tracks of each mark by taking the mean per 200bp window across the genome in hg19. We removed regions associated with repeats and blacklisted regions as given in (http://hgdownload.cse.ucsc.edu/goldenPath/hg19/encodeDCC/wgEncodeMapabilitv/wgEncodeDukeMapabilitvRegionsExcludable.bed.gz) and (http://hgdownload.cse.ucsc.edu/goldenPath/hg19/encodeDCC/wgEncodeMapabilitv/wgEncodeDacMapabilitvConsensus_Excludable.bed.gz). The processed data set contained 635 data tracks in 13.8 million windows, containing 8.8 billion observations in total. We then took log(*x*+0.1) transformation of the data as input to IDEAS.

### Robust 2D segmentation by IDEAS

Independent runs of genome segmentation may produce different results depending on the initial values of model parameters. We developed a simple but effective approach to substantially improve the reproducibility of genome segmentations between independent runs. First, we randomly selected *K* regions of 20Mb each in the genome, and ran IDEAS in each region independently. Secondly, we collected the inferred epigenetic states from the *K* runs and performed hierarchical clustering of all states based on state mean parameters. Thirdly, we identified a largest number *G* and cut the hierarchical tree of the epigenetic states into *G* or more sub trees, such that exactly *G* sub trees contained epigenetic states from at least *x*% of the *K* runs. Finally, we generated *G* consolidated epigenetic states by averaging the state parameters in each of the *G* sub trees. This approach is motivated by the following rationale: we want to identify an unknown number (*G*) of states and their parameters from multiple independent training of IDEAS; if we merge all states produced by the *K* runs together by cutting the tree at the root, we would obtain perfect reproducibility of states between runs, but with no power; on the other hand, if we treat each state from all runs as a distinct state by cutting the tree at the leaves, we would have poor reproducibility and obtain too many states; as we move down the tree from the root to the leaves, the number of sub trees will increase, so that we can find a maximum number of sub trees, within *G* of which we have states clustered together by their similarity (and hence reproducibility) from at least *x*% of the *K* runs; as we move further down the tree towards the leaves, the number of sub trees satisfying this criterion will decrease, as the total number of sub trees will increase, with fewer states in each sub tree. Using this approach, we will find a maximum number of states that satisfies the criterion that the state is reproducible in at least *x*% of the *K* runs. To improve robustness of this approach, we determined the number of states (*G*) by leave-one-out experiment, i.e., by leaving the states from each of the *K* runs out, respectively. We calculated an average number of *G* from those obtained in each leave-one-out experiment, which is then robust to outliers. Given *G*, we finally used the full tree on all states from *K* runs to obtain the consolidated states. The full tree may have more than *G* sub trees satisfying the criterion, and we simply used the results from the first feasible solution nearest to the root of the tree.

Our approach for generating reproducible states only requires the user to specify one parameter, *x*% reproducibility. If *x* is too small, we may obtain a large number of less reproducible epigenetic states. If *x* is too large, we may obtain a small number of highly reproducible states, but miss some important states. In this study, we let *x*=90, i.e., 90% reproducibility. Another parameter may be determined by the user is K, the number of independent trainings. Our procedure was not sensitive to the choice of *K* for *K*>=10, and hence we used *K*=15.

Finally, given the reproducible states identified by the above procedure, we ran IDEAS to segment the whole genome of 127 epigenomes using those state parameters as priors. To improve computational efficiency, we used parallelization. For both training and whole genome segmentation, we ran IDEAS in 20 iterations. We tried running IDEAS in 100 iterations, but the results were not significantly better than using just 20 iterations. That is, our training pipeline not only improved reproducibility but also enabled shorter runs of IDEAS.

### ChromHMM result

We downloaded the 15-state model by ChromHMM from the Roadmap Epigenomics project website (http://egg2.wustl.edu/roadmap/web_portal/chr_state_learning.html#core_15state). We mapped the ChromHMM states to the same set of windows used by IDEAS.

### RNA-seq data analysis

We downloaded RNA-seq RPKM data in 56 cell types (excluding E000) from the Roadmap Epigenomics Project (http://www.roadmapproject.org/). For each gene (Gencode.v10,Harrow et al., 2012) in each cell type, we calculated the proportions of epigenetic states in the regions from 110kb upstream (relative to the strand of the gene) of the gene’s TSS to 110kb downstream of the gene’s TTS. The proportions of states were calculated by weighted average, where weights were given by B-splines. We defined B-splines at degree 5 using 30 knots evenly spread over an [01] interval, which yielded 36 B-splines. The position 110kb upstream of TSS of a gene corresponded to 0. The position 110kb downstream of TTS of a gene corresponded to 1, and the TSS and TTS corresponded to 0.4 and 0.6 in the interval, respectively. The positions upstream of TSS were mapped to the interval [0,0.4) in log10 scale, i.e., the positions 10^k^ relative to TSS were mapped evenly between [0,0.4) with respect to *k*. Similarly, the positions downstream of TTS were mapped to the interval (0.6,1]. Finally, the positions within a gene were evenly mapped to the interval [0.4,0.6]. The weighted average of state proportions were calculated by using each B-spline as weights separately, followed by log(*x*+1e−5) transformation. The resulted values from all B-splines were used as predictors, and the RPKM values of each gene were used as responses.

We used linear regression to calculate adjusted *r^2^*, which accounted for the different degrees of freedom in the model. An overall model including the predictors from all B-splines were used to evaluate the overall prediction power of epigenetic states on expression. We also used the predictors calculated from each B-spline separately to evaluate the prediction power of epigenetic states in each region relative to TSS and TTS. To further evaluate the contribution of each epigenetic state to expression at each location relative to the gene, we calculated partial *r^2^,* i.e., by leaving each epigenetic state out, respectively. We then calculated the ratio between the partial *r^2^* of each epigenetic state and the sum of partial *r^2^* of all epigenetic states, which reflected how much each state contributed to the expression relative to the contribution by all states.

### GTEx data analysis

We downloaded the GTEx eQTLs in 44 tissues (v6p) from the GTEx Portal (http://www.gtexportal.org/home/). Within each tissue, we grouped significant eQTLs (p-val < 1e-5) within 50kb to each other together, where overlapping groups of eQTLs were further merged. We then extended the interval of each group of eQTLs to the sides by 2kb. Within each eQTL interval, we calculated a weighted state proportion, where the weight at each position was given by Σ_*i*_{–log(*p_i_*)exp(-*d_i_*)}, where *i* denotes the *i*th eQTL in the interval, *p_i_* and *d_i_* denote the nominal p-value of the *i*th eQTL and its distance to the current position. In this way, all epigenetic states within an eQTL interval were integrated, with more weights given to the positions closer to eQTLs. Since eQTLs are enriched in genic regions, instead of using genomic background, we used the same eQTL interval as controls by calculating a inversely-weighted state proportion, i.e., using inverse weights. We finally took log(*x*+1e-5) transformation of the weighted state proportions and used the values as predictors. The response variable was binary indicating cases and controls. We used logistic regression to predict eQTL intervals in each tissue by the epigenetic states in each Roadmap Epigenomics cell type respectively.

### FANTOM5 data analysis

We downloaded the CAGE based enhancer data (phase 1 and phase 2 combined) from FANTOM5 website (http://fantom.gsc.riken.ip/5/data/). There are two types of data: the tag counts of expression data normalized as tags per million mapped reads (TPM), and the binary peaks called at a significance threshold by contrasting to control data, both available in 808 human CAGE libraries. For the regression analysis, we calculated the log transformed proportions of states in each enhancer region as the predictors, regardless of whether the enhancers are active or repressed in each CAGE library, and we used the TPM values in each CAGE library as the response variable. To match between CAGE libraries and Roadmap cell types, we manually assigned each of the 808 CAGE libraries to one of the cell type categories defined by the Roadmap Epigenomics consortium. For the state composition analysis, we calculated the state proportions only within the expression peaks in each CAGE library. Finally, we downloaded the pre-calculated enhancer-TSS association data from http://enhancer.binf.ku.dk/presets/enhancer_tss_associations.bed and calculated the proportions of states within the paired enhancer-TSS regions.

### Sequence-based functional score analysis

We downloaded the GERP elements on hg19 from http://mendel.stanford.edu/SidowLab/downloads/gerp/. We downloaded the CADD score pre-calculated on 1000 Genome phase 3 variants from http://cadd.gs.washington.edu/download/, and we used the scaled score (PHRED-like score) in this study. We downloaded the FitCons score from http://compgen.cshl.edu/fitCons/0downloads/tracks/current/i6/scores/, where we used the highly significant scores (p<0.003) integrated across the three ENCODE cell types. We obtained the pre-calculated CATO scores in 13.4 million SNPs overlapping with DHS from http://www.uwencode.org/proi/CATO/. We mapped all four scores to the 200bp windows used in this study by obtaining the maximum score within each 200bp window, and we assigned 0 to the windows without scores. We transformed the scores by log((*x*+1e−4)/(1-*x*+1e−4)), which was used as the response variable. The epigenetic states in the corresponding windows were used as dummy predictors. We further shifted the window to the left and right by up to 25 windows (corresponding to 5kb) to evaluate the location precision of epigenetic states. Except for FitCons, all other scores were calculated without using cell type specific information. We thus performed regression analysis in each of the 127 Roadmap cell types separately. We further performed state enrichment analysis with respect to 50 equal-size partitions of scores, i.e., each partition has the same number of scores.

### Promoter-capture HiC data analysis

We obtained the significant promoter-interacting regions (PIR) (CHiCAGO interaction scores >5) from Data S1 of Javierre et al. (2016). We used the log(*x*+1) transformation of the CHiCAGO interaction scores in all PIRs from the 17 IHEC blood cell types as the response variable, and we used the log(*x*+1e−5) transformed state proportions in the corresponding PIRs as predictors. For each PIR, the state proportions were calculated in both the bait and the target regions, which were used as predictors in either an additive or a multiplicative way. In the latter case, the state proportions between the bait and the target regions were multiplied between all state pairs. We performed regression analysis on the PIRs in each IHEC blood cell type using states in each Roadmap cell type separately.

Blood cell type specificity was calculated based on RPKM values from the Roadmap RNA-seq data. The Roadmap blood cell types we used were E037 BLD.CD4.MPC, E038 BLD.CD4.NPC, E047 BLD.CD8.NPC, E050 BLD.MOB.CD34.PC.F, E062 BLD.PER.MONUC.PC, E123 BLD.K562.CNCR. At each gene, we calculated the mean and the variance of RPKM values in the blood cell types, as well as in non-blood cell types, all in log(*x*+1) transformed scale. The variance for each gene in each cell type group was calculated by first applying a loess smooth regression to fit the variance on the mean of genes. The variance for each gene was then taken as the value on the loess curve, or 0.25, whichever is greater, at the mean of the gene, within blood and non-blood cell types respectively. We finally calculated a two-sample t-statistic between the blood and nonblood cell types for each gene and used it as z-scores.

## ACKNOWLEDGEMENT

Zhang and Hardison are supported by grant NIH 1R24DK106766.

---

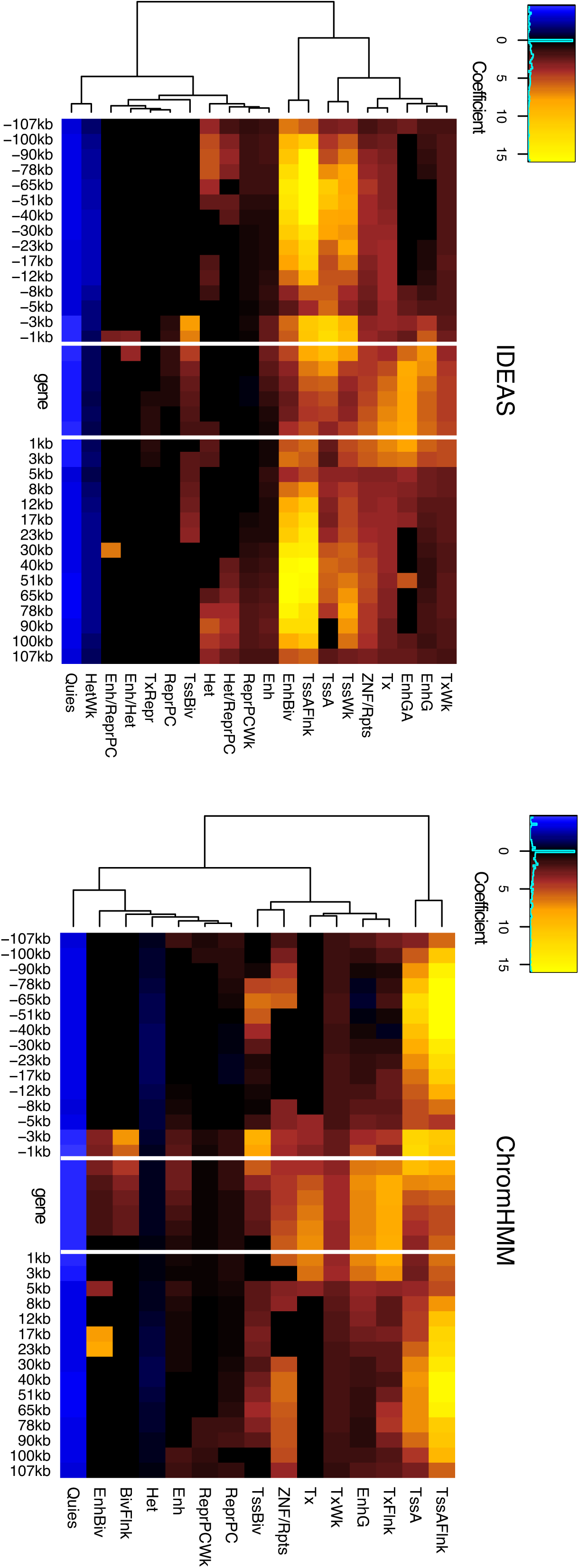

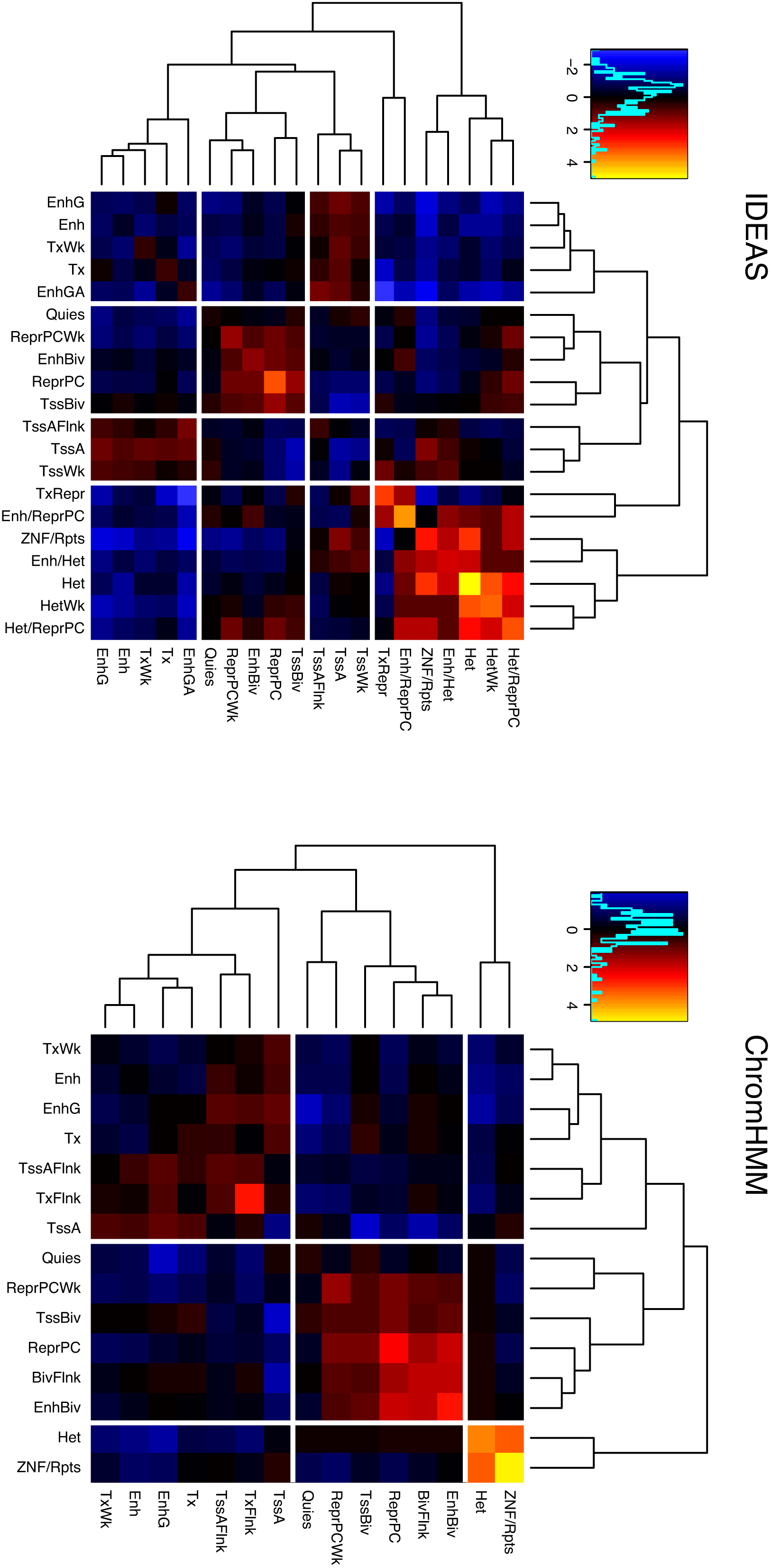

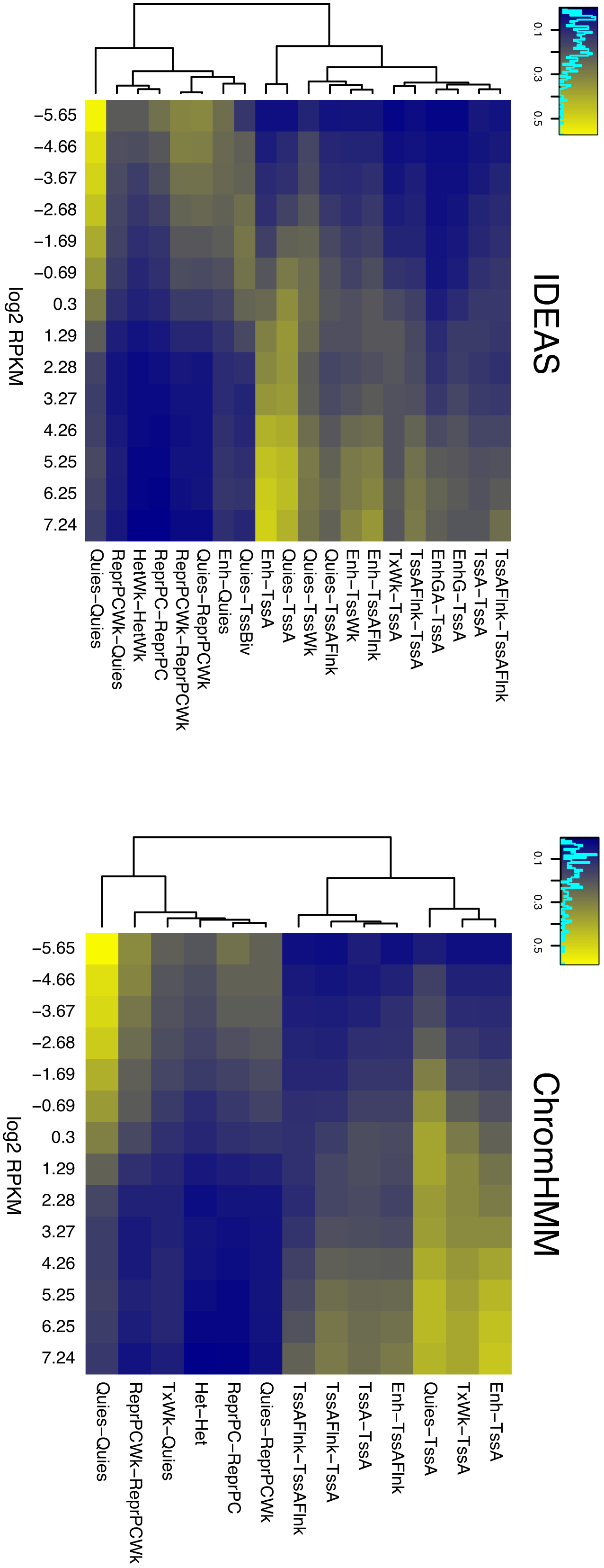

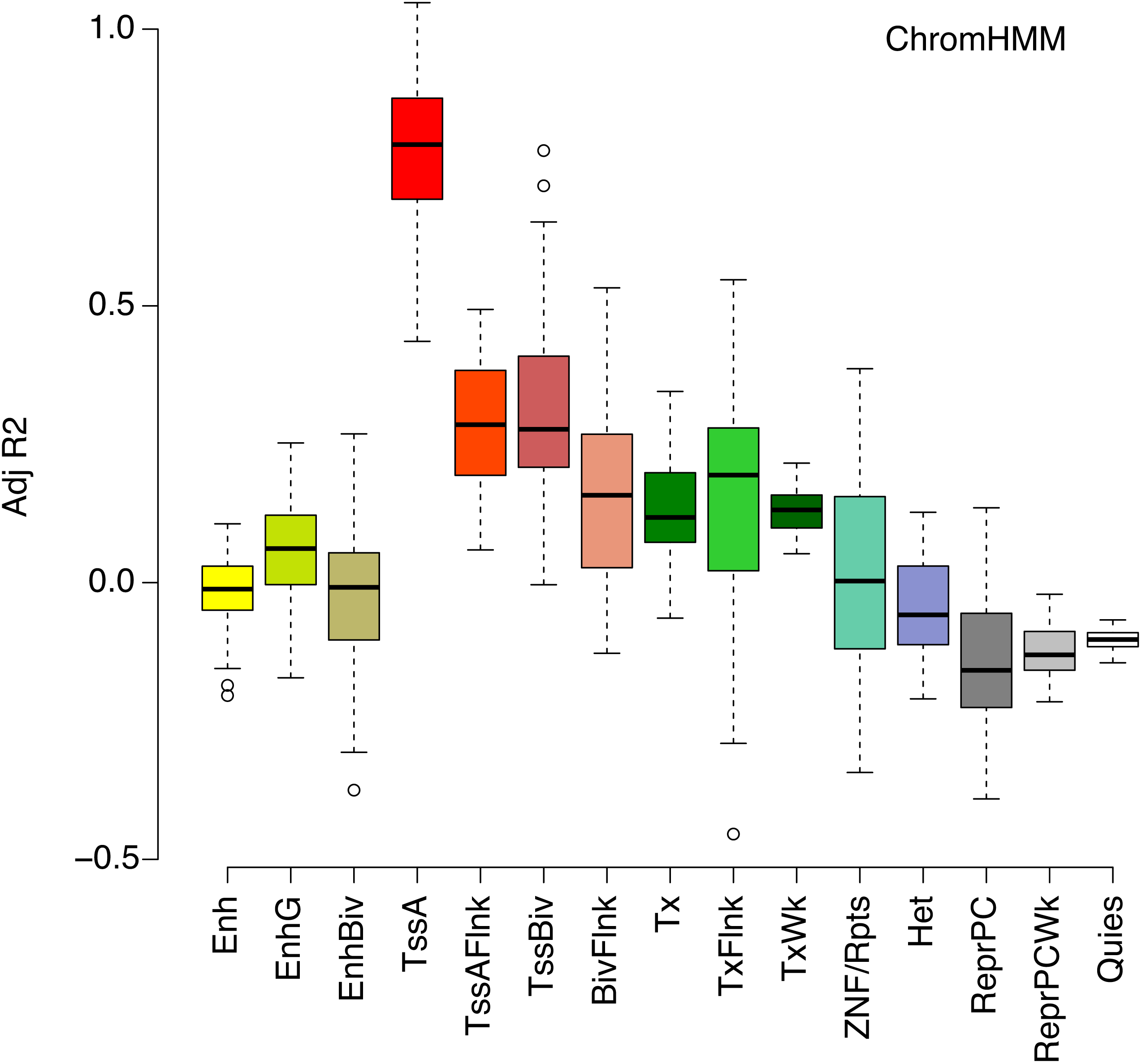

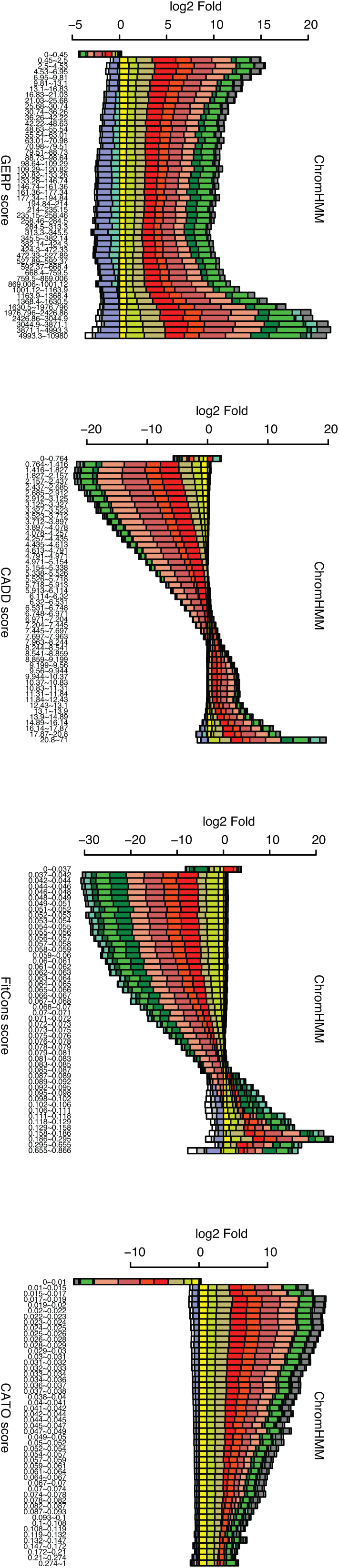

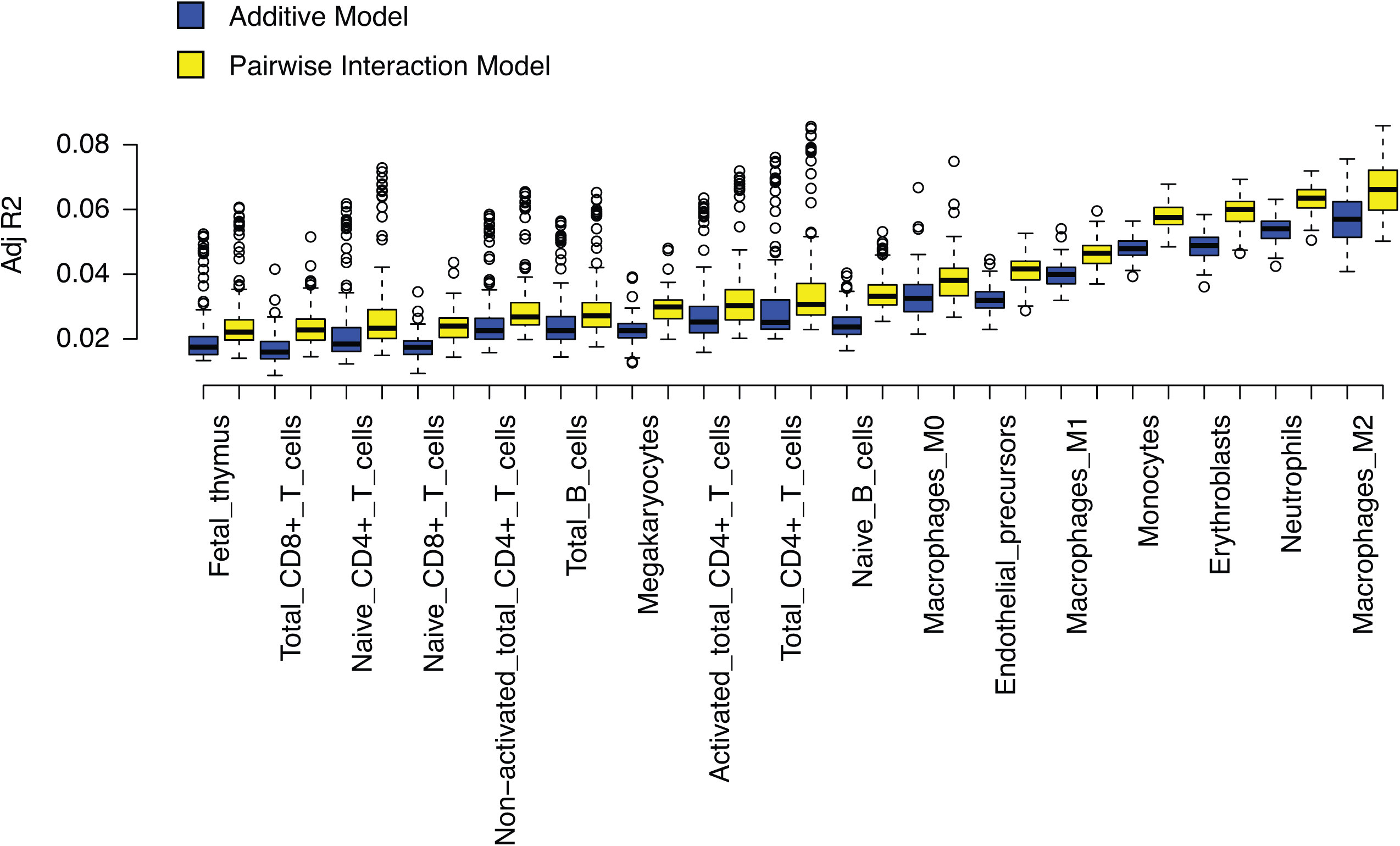

